# In vivo screening reveals new regulators of Natural Killer cell development and functional response to acute cytomegalovirus infection

**DOI:** 10.64898/2026.03.19.709783

**Authors:** EJ Pring, S Clare, R Hare, S Sheppard, M Marsden, INTREPID consortium, M Clement, L Chapman, K Harcourt, C. Ballesteros Reviriego, C Brandt, A Andres, B Abu-Helil, V Iyer, L van der Weyden, MA Oliver, PA Lyons, ECY Wang, DJ Adams, MC Cook, DM Davis, IR Humphreys, AO Speak

## Abstract

Natural killer (NK) cells are critical innate immune effectors in antiviral and antitumour responses. However, the factors governing NK cell development and function remain incompletely understood. To address this, we adopted an *in vivo* screening approach, generating a panel of mice deficient in genes differentially expressed across NK cell maturation stages or following viral infection. We performed a combinational screen comprising baseline immunophenotyping and mouse cytomegalovirus (MCMV) challenge. We identified novel regulators of NK cell development, including transcriptional regulators and proteins with putative trafficking functions. MCMV challenge studies additionally identified proteins that impacted antiviral immunity independently of NK cell phenotypes, including a novel regulator of NK cell degranulation, Synaptotagmin-like protein 3. Identified genes exhibited reduced tolerance to loss-of-function in large scale human sequencing studies and evidence for reduced NK cell numbers in a cohort of immunodeficiency patients. Together, these findings provide a resource of NK-expressed genes important for NK cell maturation and function, with relevance to human health.

**Summary:** Pring et al. conduct an *in vivo* screen in mice identifying novel regulators of natural killer (NK) cell development and/or antiviral functionality. Large-scale human sequencing and human immunodeficiency patients revealed evidence for the relevance of identified genes in human health.

## Introduction

Natural killer (NK) cells represent a vital first line of defence against viral infections and have the capability to recognize and kill transformed cells. Control of NK cell activation is achieved by the integration of inhibitory signals, such as those provided by major histocompatibility complex I (MHC class I) on the surface of target cells (‘self’) and activating signals, for example the NKG2D ligands MICA, MICB and ULBP family members ^1^. The required threshold for NK cell activation is determined by the balance of these inhibitory and activating signals and can be further tuned by cytokines ^2^. Although it is clear that NK cell development is dependent on IL-15 and requires the action of transcription factors such as *Tbx21*, *Eomes*, *Ets1*, *Nfil3*, *Id2* and *Tox1* (reviewed in ^3^), other factors that determine NK cell development and/or the associated expression of receptors that may determine NK activation thresholds are unknown.

NK cell development in mice can be divided into four stages through surface expression of CD27 and CD11b^4^. The function of NK cells through these stages differs, with CD27^hi^ CD11b^hi^ NK cells exhibiting elevated cytotoxicity and cytokine secretion after stimulation as compared to CD27^lo^ CD11b^hi^ NK cells ^5^. This correlates with differential expression of stimulatory and inhibitory receptors. In addition, these subsets possess differential proliferative capacity and migration towards chemokines and sphingosine-1-phosphate ^5,6^. Previous studies have investigated the pathways of NK cell maturation through CD27^hi^ CD11b^lo^ to CD27^lo^ CD11b^hi^ at a molecular level via microarray profiling of gene expression^4^. In addition, NK cell gene expression in response to viral infection, using the mouse cytomegalovirus (MCMV) model, has also been investigated ^7^.

We hypothesized that differential expression of NK cell genes during development and/or in response to viral infection may reveal important unknown determinants of NK cell function and development. Thus, we generated a panel of gene-deficient mice selected from the most differentially regulated genes during NK cell development ^4^ or in response to viral infection ^7^. These mice were subjected to a broad immunophenotyping panel to define any baseline phenotypes, and their susceptibility to viral infection was assessed using the MCMV model. Using this *in vivo* approach, we identified a number of genes that impact NK cell number, development and/or antiviral function. Furthermore, genes identified in our mouse screen were identified as more intolerant to loss of function in humans, and variants in these genes were identified in a cohort of primary immunodeficiency patients with deficiencies some presenting with reduced circulating NK cells.

## Results and Discussion

### Selection of candidate genes for in vivo screen

To identify candidate genes for *in vivo* screening, we combined existing microarray datasets ^4,7^ and a single cell dataset from index sorted NK cells isolated from wild type (WT) C57BL6/N mice. The combined gene set was then filtered to remove genes where knockout mice had previously been generated and then triaged for the ability to design guides for CRISPR-Cas9 mediated gene deletion. A total of 67 gene targeted mouse lines were successfully generated including a triple-targeted allele to delete the whole *Klrc1-3* region. We also incorporated a EUCOM/KOMP ES cell-generated *Zfp292*-deficient mouse line.

To validate and improve the resolution of previously generated microarray data ^4^, we performed bulk RNAseq on isolated CD27^hi^ CD11b^lo^ and CD27^lo^ CD11b^hi^ NK cells from the spleens of C57BL6/N mice. We identified 2954 protein-coding genes that were differentially expressed between these cell subsets (DEseq2 FDR 5%, Fig. S1) including 217 of the 357 genes (60.6% recovery) previously identified in microarray studies. Furthermore, we identified that 10 genes selected from the MCMV infection microarray data were also differentially expressed during NK development (upregulated during maturation: *Dnajb9*, *Far1, Gcc2, Ttc14* and *Zfp292*; downregulated during maturation: *Ccda2, Klra6, Mrpl20, Ncapd2* and *Palm*). These data highlight that the increased depth afforded by RNAseq yields a more complete understanding of gene expression differences during NK cell development.

### Transcriptional regulators and trafficking proteins regulate mouse NK cell development

The 67 mutant alleles were expanded to generate homozygote mice, and 8 lines were found to be lethal (no homozygote animals recovered from ≥ 28 P14 offspring of heterozygote x heterozygote intercrosses). Another 6 lines were subviable (≤ 13% P14 offspring of heterozygote x heterozygote intercrosses) so both heterozygous and homozygous mice were investigated for some of these cases. While *Ccdc85c* homozygote offspring were viable, they presented with severe hydrocephalus and thus only heterozygote mice were tested. Targeting *Klrc3* yielded homozygote male mice that were often misidentified as female mice due to the abnormal external genital appearance. This was not observed in single targeting of *Klrc1* or *Klrc2*, nor the triple knockout (KO) of the whole region, the reasons underpinning this observation were unclear.

A total of 5 female and 5 male mutant mice for each line were analysed using two flow cytometry panels on spleen and blood to enable identification of the major immune cell populations and NK cell developmental/function markers (CD27, CD11b, KLRG1 and Ly49H). Tissues were analysed from mice aged 12-16 weeks of age together with age- and gender-matched WT controls. A standardised flow cytometer and gating strategy was used to analyse the frequencies of cell populations in a blinded manner by a single operator. Statistical analysis was performed using the PhenStat package ^8^ with the time as a fixed effect test to account for the multi batch workflow and P values were corrected over the whole data set for each tissue (Benjamani-Hochberg correction). Phenodeviants were identified by selecting those with an adjusted P value of < 0.05 and also using an effect size threshold (Cohen’s *d* 0.8) to prioritise those with stronger phenotypes. As the phenotyping panel included B and T cell markers there was a broad range of immunophenotypes observed which included pleiotropic effects across multiple cell subsets and tissues to very discrete phenotypes (all data from the immunophenotyping is publicly available on Zendo DOI 10.5281/zenodo.18879757).

For NK cell parameters, genes could be broadly divided into three main categories where a significant difference was observed in both sexes. Genes affecting both the abundance and phenotypic markers such as *Zfp292, Chft8, Gcc2, Scaf1 and Ergic3*; those that only affected the percentage of NK cells such as *Dnajc1* and *Glcci1;* and finally gene deletions that only affected surface marker expression such as *Etv3, Dnajb9, Trerf1* and *Zfp870* knockout mice (Fig. 1 a).

**Figure 1.**
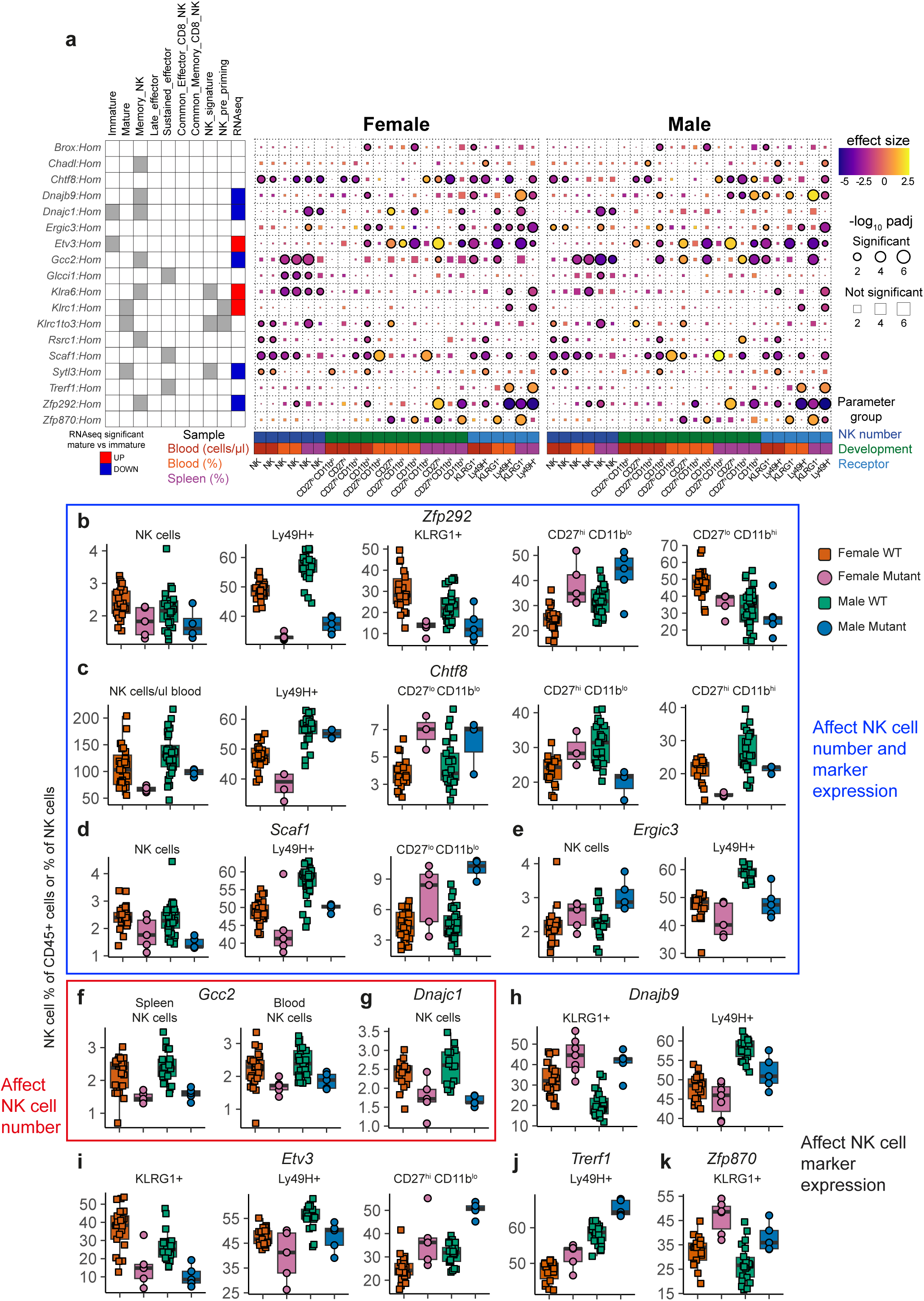
Identification of NK cell phenotypes. **(a)** Heatmap of significant NK related parameters from high throughput immunophenotyping screen. Left hand side indicates gene assignment (grey shading is present) from previous microarray experiments immature and mature NK ^4^ or effector functions ^7^. RNAseq is coloured blue for genes significantly down regulated and red significantly up in immature versus mature NK cells (from Fig. S1). Hom = homozygote and Het = heterozygote for the genotypes. **(b-k)** Example data from spleen (or blood where indicated) immunophenotyping. NK cells are expressed as % of CD45+ live cells from the tissue and NK phenotypic subsets are % of NK cells or as cells/μl blood. Each symbol represents an individual animal with the gene and parameter listed above the plot, all mutant mice are homozygote. All reported are significant findings with padj < 0.05 and effect size > Cohen’s d 0.8 and significant in both sexes, full statistical analysis and data are available on Zendo 10.5281/zenodo.18879757.

*Zfp292* is a zinc finger transcription factor that regulates innate lymphoid cell (ILC) type 3 maintenance ^9^. Mice lacking *Zfp292* exhibited reduced frequencies of circulating NK cells, with those detectable exhibiting an immature (CD27^hi^ CD11b^lo^) phenotype and reduced expression of KLRG1 and the activating receptor Ly49H (Fig. 1 b). Consistent with a reduced maturation phenotype, *Zfp292*^−/−^ mice revealed impaired *in vivo* clearance of MHC-class I deficient target cells (Fig. S2 a). *Chtf8* forms part of the Ctf18 replication factor C complex involved in sister chromatid cohesion and highly conserved from yeast ^10^. *Chtf8*^−/−^mice have reduced circulating NK cells, reduced expression of Ly49H as well as developmental defects correlating with increased immature CD27^lo^ CD11b^lo^ and a reduction of the more mature CD27^hi^ CD11b^hi^ subset (Fig. 1 c). *Scaf1* may function in pre-mRNA splicing and in pancreatic cancer regulates the deubiquitinating enzyme USP15 ^11^, which is a key regulator of numerous immune cell functions ^12^. *Scaf1^−/−^* mice exhibited reduced NK cell frequencies, Ly49H expression (Fig. 1 d) and impaired clearance of MHC-class I-deficient target cells (Fig. S2 b). *Ergic3* encodes a trafficking protein that regulates ER-to-Golgi transport of gap proteins^13^. *Ergic3*^−/−^ mice displayed increased NK cell frequencies and decreased Ly49H expression (Fig. 1 e).

Another trafficking protein that regulated NK cell accumulation was the Golgi resident coiled coil protein Gcc2 (Fig. 1 f). Two J protein family members, *Dnajc1* and *Dnajb9,* that act as co-chaperones to HSP70 ^14^ impacted NK cell development. *Dnajc1* knockout reduced NK cell frequencies (Fig. 1 g) whereas *Dnajb9^−/−^*mice exhibited altered surface marker/receptor expression (Fig. 1 h). The transcription factor *Etv3* alters monocyte differentiation by repressing macrophage commitment to favour dendritic cell generation^15^ but a role in lymphocytes has not previously been ascribed. Here we observed numerous phenotypes including increased monocyte and dendritic cells in line with the previous report ^15^ as well as an elevated B cell percentage with reduced αβ T cells (Fig. S2 c). The phenotype of the αβ T cells was altered with increased CD25^hi^ regulatory CD4+ αβ T cells and CD44^hi^ CD62L^lo^ (Fig. S2 c). There were also effects on NK cell maturation with an increased fraction of CD27^hi^ CD11b^lo^ NK cells and decreased KLRG1 and Ly49H expression (Fig. 1 i). There are also genes that seem to affect expression of a single marker on NK cells including *Trerf1* whose knockout led to increased Ly49H^+^ NK cells (Fig. 1 j) and *Zfp870* which had increased KLRG1^+^ NK cells (Fig. 1 k). These are not exhaustive lists, and all could represent novel genes that affect immune cell biology and new areas for further investigation. However, these data represent clear evidence of roles for novel transcription factors and other transcriptional regulators, and trafficking proteins as key regulators of NK cell development.

### Identification of antiviral effector genes using an in vivo MCMV infection screen

In addition to baseline immunophenotyping, all mutant alleles were subjected to MCMV challenge to identify genes that contribute to antiviral immune control. MCMV is a powerful tool for identifying mechanisms that govern antiviral NK cell responses. C57BL6 mice exhibit genetic resistance to MCMV due to expression of Ly49H which specifically binds the MCMV glycoprotein m157 on the surface of infected cells ^16,17^. Upon activation by MCMV, a clonal expansion and retention of a pool of Ly49H^+^ memory-like NK cells is induced ^18–20^. NK cell-mediated control of MCMV replication *in vivo* is well-defined, with a predominant role for perforin-dependent and Ly49H-mediated control of MCMV replication in the spleen in C57BL6 mice ^19,21,22^. For subviable lines where the single cohort of homozygote mice needed for infections could not be generated, heterozygote mice were tested. After infection, weight loss was monitored daily and after four days viral load in the spleen and liver was determined (Fig. 2 a).

**Figure 2.**
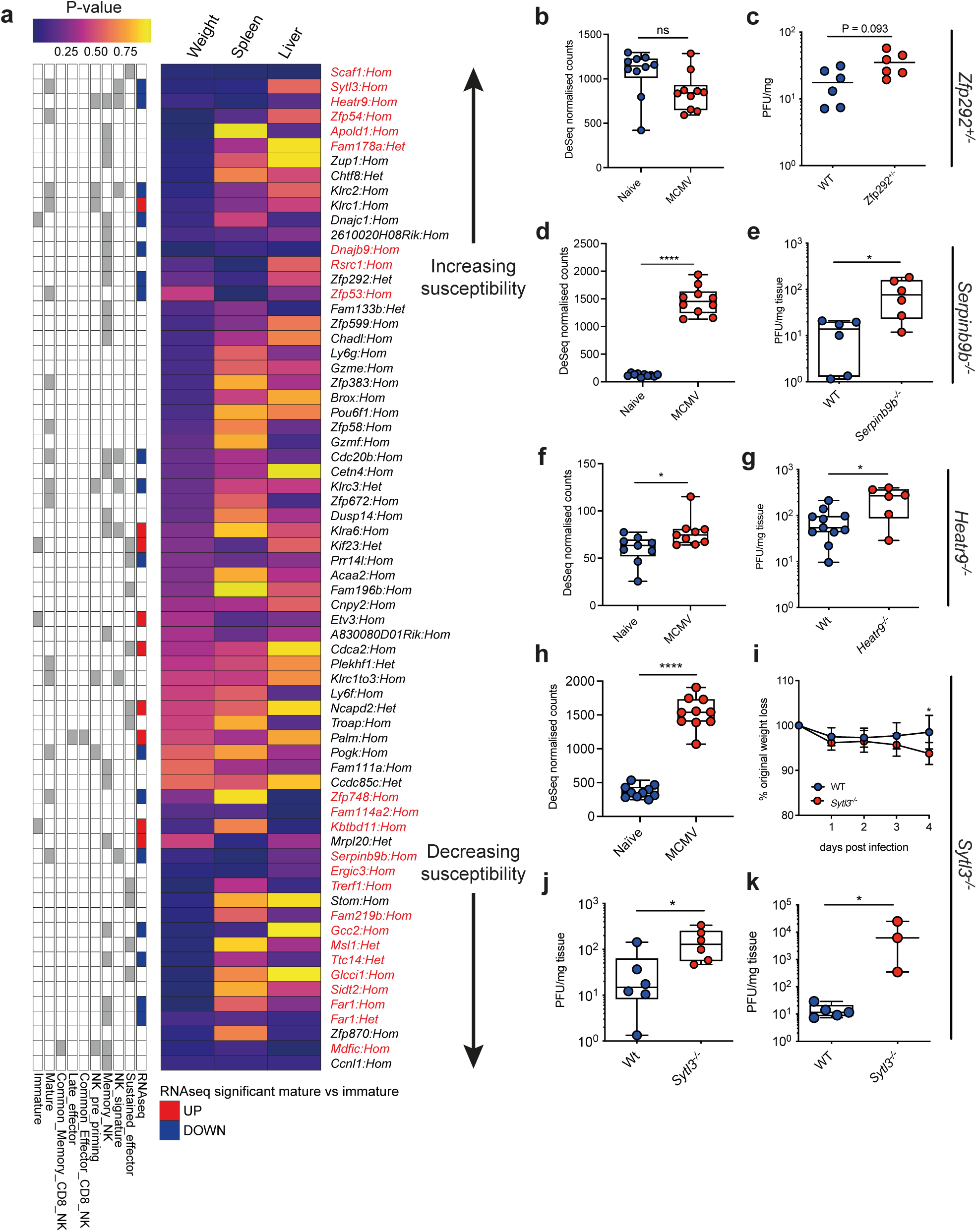
MCMV screen identifies multiple regulators of antiviral immunity. **(a)** Heatmap of p value from MCMV screen sorted according to effect on virus-induced weight change, and virus control in spleen and liver (as indicated by arrows), genes in red were significant in the screen (p < 0.05). Left hand side indicates gene assignment (grey shading is present) from previous microarray experiments immature and mature NK ^4^ or effector functions ^7^. RNAseq is coloured blue for genes significantly down regulated and red significantly up in immature versus mature NK cells (from Fig. S1). Hom = homozygote and Het = heterozygote for the genotypes **(b, d, f, h)** Gene expression of *Zfp292 **(b)**, Heatr9* **(d)**, *Serpinb9b* **(f)**, or *Sytl3* **(h)** in blood of naïve mice and 2 days post-MCMV infection (from ^48^). Individual mice (10/group) + range are shown. **(c, e, g)** MCMV titers 4 days post infection in spleens of WT and *Zfp292^+/−^ **(c),** Heatr9^−/−^* **(e)** or *Serpinnb9b^−/−^* **(g)** mice was assessed by plaque assay. Weight loss **(i)** and MCMV titers 4 days post infection in spleens **(j & k)** of WT and *Sytl3^−/−^* mice was quantified by plaque assay following infection with tissue culture-derived **(i & j)** and salivary gland-derived **(k)** MCMV. Individual mice are shown, and data represent > 4 **(i & j)** or 2 **(k)** experiments. Statistical analysis was performed using the Mann-Whitney U test. * p < 0.05, **** p < 0.0001.

A total of 23 genes were classified as affecting the response to MCMV with enhanced and impaired responders identified based on virus load in spleen and virus-induced weight alterations (Fig. 2 a). Notably, altered NK cell development and/or receptor expression did not always predict impaired antiviral control, as evidenced by unaltered and increased protection from MCMV infection in *Etv3^−/−^* and *Dnajb9*^−/−^ mice, respectively (Fig. 2 a). However, although *Zfp292* expression in peripheral blood was not induced by MCMV infection (Fig. 2 b), heterozygote mice indicated a trend in increased virus replication in the spleen (Fig. 2 c). Although the sub-viable nature of *Zfp292^−/−^* precluded investigation of homozygote mice, these data implied that impaired NK development due to *Zfp292* deficiency impacts antiviral control.

We also identified multiple genes that limited MCMV pathogenesis without impacting NK development. Indeed, many genes that afforded protection from MCMV pathogenesis (virus replication and/or weight loss) were selected based on increased expression by mature NK cells, consistent with the enriched expression of known antiviral effector proteins such as granzyme B and FasL by mature NK cells ^4^. Examples included *Serpinb9b*, which encodes a protein orthologous to human serine protease inhibitor SERPINB9, which is a known inhibitor of granzymes ^23^. It is broadly expressed by hematopoietic and non-hematopoietic cells ^24^ but in the context of NK cells, promotes survival of granzyme B^+^ NK cells and host protection from poxvirus infection ^25^. MCMV induced robust *Serpinb9b* expression in peripheral blood (Fig. 2 d) and, although no impact of gene deficiency on weight loss was observed, *Serpinb9b* knockout mice exhibited increased splenic virus replication 4 days-post infection (Fig. 2 e). Novel antiviral effector genes that are preferentially expressed by memory NK cells were also identified. For example, *Heatr9*, which is conserved in mice and humans but has no known function, was induced by MCMV infection (Fig. 2 f) including by memory NK cells ^7^ and restricted splenic MCMV replication (Fig. 2 g).

The top-ranking gene in our screen that impacted MCMV control and weight loss was Sytl3 (Synaptotagmin-like protein 3). The *Sytl* gene family encodes proteins with a similar structure to synaptotagmins which have an established role in vesicular trafficking events including mast cell exocytosis ^26^ and macrophage phagocytosis ^27^. *Sytl3* is predominantly expressed by NK cells ^7^, basophils and, to a lesser extent, T cells ^28^, and was identified to be up-regulated by mature (CD27^lo^) NK cells ^4^. *Sytl3* expression was markedly elevated in the blood of MCMV-infected mice (Fig. 2 h) and in our screen *Sytl3^−/−^* mice were one of the few lines that exhibited a statistically significant increase in virus-induced weight loss (Fig. 2 i). Elevated virus load was observed in the spleens of *Sytl3^−/−^* mice infected with our standard tissue culture-derived (Fig. 2 j) and when challenged with the more pathogenic salivary gland-derived MCMV (Fig. 2 k).

### Sytl3 is a determinant of NK cell degranulation

We sought to further understand the importance of *Sytl3* in antiviral NK cell responses. Splenic viral loads were comparable in NK-depleted WT and *Sytl3^−/−^* mice, implying that defective NK-mediated control of viral replication contributed to the susceptibility of *Sytl3^−/−^*mice to MCMV infection (Fig. 3 a). Despite the enrichment of Sytl3 expression in mature NK cells, naïve *Sytl3^−/−^* mice exhibited unaltered NK cell abundance, development and activating receptor expression (Fig. S3 a-c). Upon MCMV challenge, *Sytl3^−/−^* mice also exhibited no defects in NK cell accumulation, developmental subsets, expression of Ly49H and the marker of activation CD25, or granzyme B (Fig. S3 d-g).

**Figure 3.**
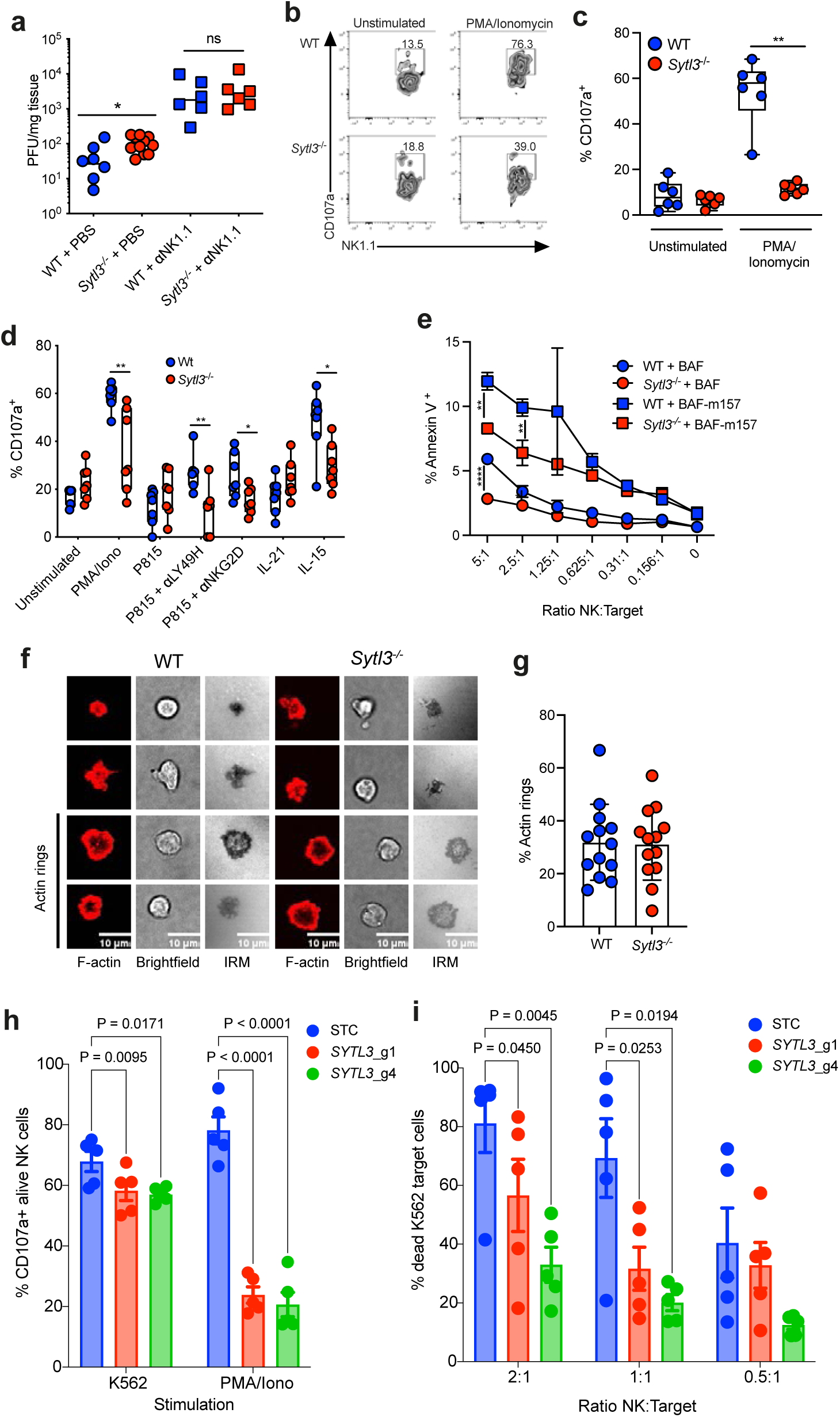
Syt3 promotes NK cell cytotoxicity in response to multiple stimuli in a cell-intrinsic manner. **(a)** MCMV titers in the spleen 4 days after infection with tissue culture-derived MCMV +/− anti-NK1.1 administration. Data are merged from two experiments and represent 4 in total. **(b & c)** Representative FACS plots **(b)** and quantification **(c)** of degranulating NK cells from spleens of WT and *Sytl3^−/−^* mice 4 days after MCMV infection. Data represent > 3 experiments. **(d)** Externalization of CD107a was assessed by flow cytometry following stimulation of NK cells from WT and *Sytl3*^−/−^ mice +/− PMA-Ionomycin, cytokines or P815 cells pre-incubated +/− agonist antibodies to activating NK cell receptors. Cells were pooled within groups where lymphocyte numbers were insufficient. Individual mice (except 1 point which represents 2 pooled mouse samples), median and interquartile range are shown. Data from 1 of 3 experiments are shown. Significance was assessed using paired analysis of WT and *Sytl3*^−/−^ NK cells in each treatment condition using a Mann Whitney U test. *p < 0.05, ** p = 0.005. **(e)** NK cells derived from WT and *Sytl3^−/−^* mice were stimulated with BaF/3 control and BaF/3-m157 cells across several E:T ratios. Annexin V binding to target cells was assessed. Mean +/− SD is shown from 3 technical replicates/data point. Paired statistical analyses of WT vs *Sytl3*^−/−^ was performed using Student’s t-Test at each E:T ratio for either m157^+^ or m157^−^ cell lines. *p < 0.05. **(f)** Representative microscopy images of F-actin (red), brightfield and interference reflection microscopy [IRM] (shown in grayscale) in naïve WT and *Sytl3^−/−^*NK cells stimulated for 10 minutes on surfaces coated with anti-NK1.1 mAb. Scale bars, 10 µm. **(g)** The proportion of cells that have formed an actin ring. Data represents 2 experiments. **(h)** Degranulation of human primary blood expanded NK cells in response to K562 or PMA/Ionomycin stimulation and **(i)** specific killing of K562 targets after 3-hour incubation with NK cells. STC is a safe cutting control and SYLT3_g1 and SYLT3_g4 are two independent sgRNA to generate SYLT3 KO. **(h-i)** Each dot represents an individual human donor, p values from 2-way ANOVA with correction for multiple testing with a post-hoc Dunnett test. Representative image from two independent experiments with unique donors.

Although Sytl3 deficiency did not impact NK accumulation or phenotype, NK cells isolated from MCMV-infected *Sytl3^−/−^* mice exhibited a marked reduction in *ex-vivo* degranulation following PMA and Ionomycin stimulation (Fig. 3 b-c). In contrast, we observed no reproducible impact of *Sytl3* deficiency on IFNγ-expressing NK cell responses *ex-vivo* (data not shown). Given that NK cell cytotoxicity is the predominant effector function that controls MCMV replication in the spleen in this model ^21^, these data suggest that impaired antiviral control by *Sytl3*^−/−^ mice reflected reduced NK cell cytotoxic function rather than an impact on NK cell development, activation, survival and/or cytokine production.

We next determined whether *Sytl3* directly impacted NK cell degranulation. Purified WT and *Sytl3*^−/−^ NK cells were stimulated *in vitro* with various stimuli and NK cell degranulation was measured using flow cytometry. *Sytl3^−/−^* NK cells exhibited reduced degranulation following stimulation with PMA and Ionomycin, IL-15, and target cells bound with anti-Ly49H or anti-NKG2D antibody (Fig. 3 d). In accordance, *Sytl3^−/−^* NK cells exhibited defective killing of m157-expressing target cells (Fig. 3 e). Sytl3 deficiencies suppression of degranulation did not result from an impact on F-actin remodelling and ring formation (Fig. 3 f-g). The presence of Sytl3 (Slp3) in endosomal exocytotic vesicles of cytotoxic T cells in complex with Rab27a and kinesin-1 has been reported, and this may facilitate anterograde transport of lytic granules from the MTOC towards the immunological synapse ^29^. Moreover, related proteins Slp1 and Slp2 are implicated in the tethering of Rab27a to the T cell plasma membrane ^30,31^ (although Sytl3/Slp3 was not detected in these experiments ^31^). Our data are consistent with a role for Sytl3 in the final stages of granule exocytosis and suggest perturbations in this process impairs control of MCMV infection. Finally, we generated *SYLT3^−/−^*primary human NK cells (Fig. S3 h) and demonstrated reduced NK cell degranulation in response to PMA and Ionomycin and, to a lesser extent, K562 cells in the absence of *SYLT3* (Fig. 3 h). This correlated with reduced killing of K562 targets cells (Fig. 3 i), demonstrating that SYLT3 promotes NK degranulation and target cell killing in mice and humans.

### Relevance of NK-related genes to human biology

Given the shared role of SYTL3 in both mouse and human NK cells, we hypothesised that phenodeviant genes identified in our mouse screen could be important for human health and would be more intolerant to loss of function in humans. Using the large-scale sequencing data from GnomAD, we identified the human orthologue for the mouse genes and compared the LOEUF value ^32^. Of the genes from our mouse screen, we compared 1) genes with no phenotype, 2) genes with a phenotype only in MCMV infection (immune functional defect), 3) genes that altered NK cell parameters only and 4) genes that influenced response to MCMV and NK cell parameters (Fig. 4 a). Genes that both impacted the baseline NK phenotype and altered response to MCMV demonstrated an increased prevalence of the human orthologue having a lower LOEUF value as compared to genes without a phenotype, indicating they are more intolerant to loss in the human population (Fig. 4 a).

**Figure 4.**
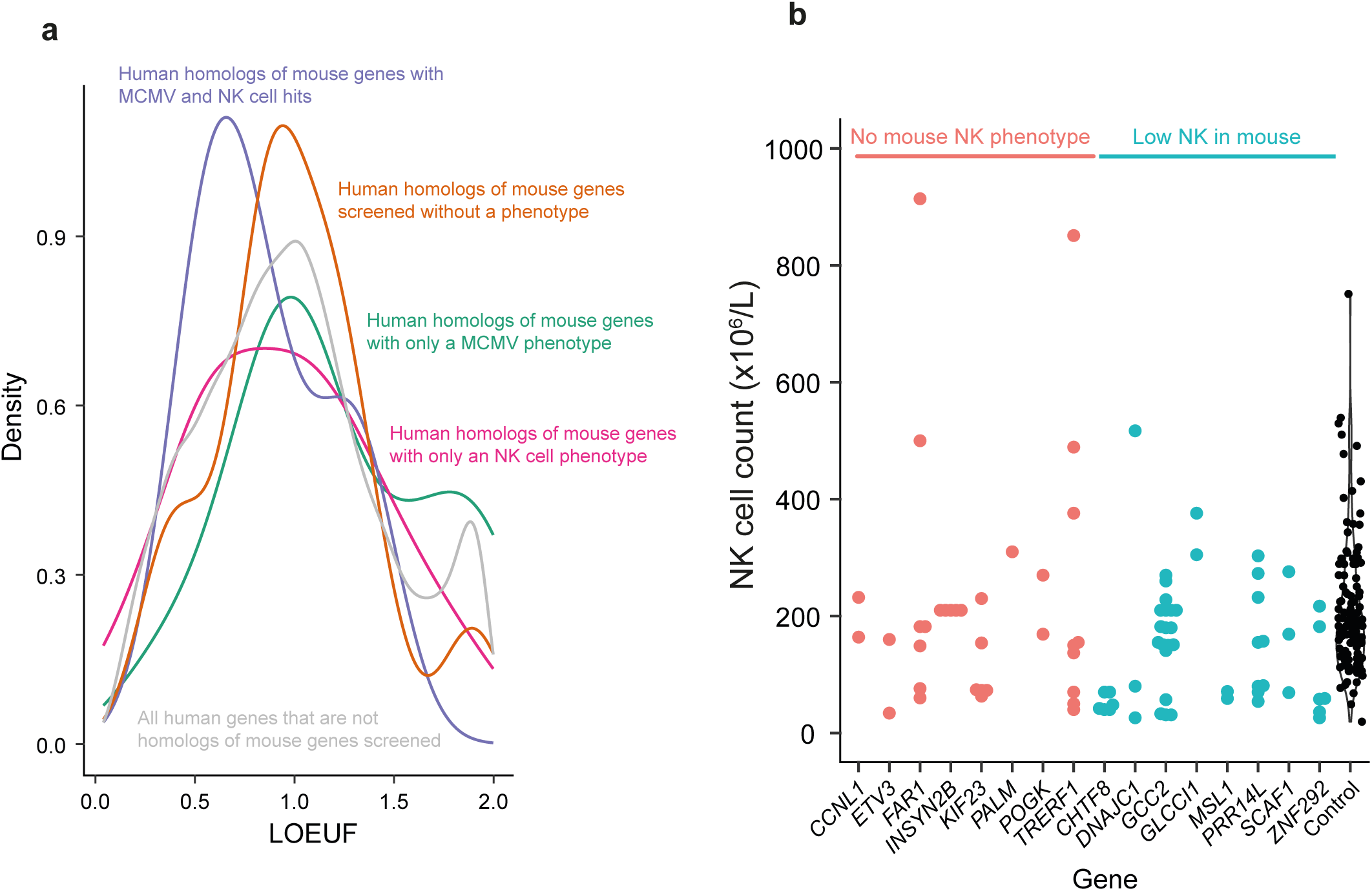
Evidence of human relevance for genes. **(a)** Density plot for LOUEF values for human orthologues of genes categorised as indicated. **(b)** NK cell counts from INTREPID cohort filtered for autosomal dominant variants in genes identified in this screen. Variants are categorised as those without a low NK cell phenotype in the mouse or those with low NK cells in the mouse. Controls are healthy donors processed alongside the patient samples. Each symbol represents an individual.

We next investigated whether any genes identified to affect NK cell numbers in mice could be identified in a large cohort of immunodeficiency patients ^33^. We filtered the cohort for the homologues of the genes and categorised the variants as those predicted to be autosomal dominant or recessive. Selecting the predicted dominant variants, we categorised the genes into those with a phenotype of low NK cells in the mouse immunophenotyping screen or those without a low NK cell phenotype and compared the NK cell count in peripheral blood (Fig. 4 b). While the numbers of individuals are quite low there are several genes that reach significance when compared to controls including *CHTF8* (P < 0.0001, Wilcox test), *ZNF292* (P = 0.0121, Wilcox test) and *MSL1* (P = 0.211, Wilcox test). *Chtf8^−/−^* mice showed reduced NK cells as well as developmental abnormalities with an increased proportion of more immature NK cells (Fig. 1 b). Apart from a role in sister chromatin cohesion ^10^ there is no indication of how this gene might affect NK cell development. *Zfp292^−/−^* (mouse homologue of *ZNF292*) also demonstrated reduced NK cell numbers and impaired development (Fig. 1 b). *Zfp292* is a transcription factor required for the maintenance of ILC3s ^9^, and given the shared early developmental origin of NK cells with other ILCs ^34^, it is interesting to speculate a similar role for *Zfp292* in NK cells with other ILC lineages. *ZNF292* is intolerant to loss of function (gnomAD LoF o/e = 0.07 (0.04-0.14)) and variants have been linked to neurodevelopmental disorders ^35^. Both *Zfp292* and *Chtf8* homozygous mice were subviable and thus it was not possible to robustly test in the MCMV infection screen, although there was a trend to impaired virus clearance in *Zfp292^+/−^* mice (Fig. 2 c). *MSL1* is part of the male-specific lethal (MSL) acetyltransferase complex responsible for adding the epigenetic H4K16ac mark ^36^ and has been shown to be important for maintaining chromosome integrity in cancer cell lines ^37^. Although it is clear that there are substantial differences in chromatin accessibility between NK and ILC subsets ^38^ there is no data regarding the role of the H4K16ac mark in NK cells. *Msl1^−/−^* mice were lethal and thus only heterozygous mice were tested, and these exhibited a reduction in circulating NK cells (DOI 10.5281/zenodo.18879757).

Thus, our mouse screen identified genes differentially expressed during NK cell development and/or antiviral responses, that lead to the discovery of key proteins that impact the development and function of human NK cells. Moreover, our study revealed that combined screening of baseline immunophenotyping and *in vivo* antiviral responsiveness in mice can be used to predict genes with a significant impact on human health. This demonstrates the power of linking mouse knockout data to clinical annotation to inform the prioritisation of immune regulating genes.

## Materials and methods

### Experimental design

All experiments performed on mice were not blinded to genotype due to this information being present on cage cards. Experiments were performed in a randomised manner on a cage basis with controls and mutant mice generally being co-housed. A multi-batch sample collection was performed for the baseline flow cytometric analysis to ensure reproducibility over multiple litters and days of phenotypes. MCMV infections were performed in a single cohort of animals. All analysis of flow cytometry data was performed in a blinded manner during sample preparation and analysis. All plaque assay experiments were performed in a blinded manner by different researchers in a different institute from where the *in vivo* experiments were performed. No *a priori* estimates were performed to calculate sample sizes for experiments and mice were allocated to treatment groups by random allocation via Mendelian inheritance. The care and use of all mice in this study were in accordance with the Home Office guidelines of the UK and procedures were performed Home Office Project Licenses (P2E57E159, P7867DADD, P96810DE8 and P653704A5) which were reviewed and approved by the Wellcome Trust Sanger Institute or Cardiff University Animal Welfare and Ethical Review Body.

### Generation of mouse lines

65 strains of gene deletion mice were generated on a C57BL6/N background by the Wellcome Trust Sanger Institute (WSI) using CRISPR-Cas9 ^39^, one large deletion spanning genes *Klrc1, Klrc2 and Klrc3* was also generated using CRISPR-Cas9 and one allele was generated using the EUCOMM/KOMP embryonic stem cell collection ^40^ as part of the Sanger Institute Mouse Genetics Project. WT C57BL6/N controls were bred in-house and typically sourced from the offspring of genetically altered lines. After founders were generated and sequenced (for CRISPR-Cas9 lines), heterozygote mice were paired to generate homozygote offspring. A number of lines were lethal (no homozygote animals recovered from ≥ 28 P14 offspring of heterozygote x heterozygote intercrosses) or subviable (≤13% P14 offspring of heterozygote x heterozygote intercrosses). For lethal lines, heterozygote mice were analysed and for subviable lines, either heterozygote, homozygote or both genotypes were used depending on breeding performance and viability.

### RNA Sequencing

Single cell suspensions were prepared from spleens obtained from WT mice and stained with titrated cocktail of fluorochrome conjugated antibodies (anti-NK1.1 (clone PK136), anti-CD3 (clone 145-2C11), anti-CD27 (clone LG.3A10), anti-CD11b (clone M1/70) and anti-KLRG1 (clone 2F1/KLRG1) all Biolegend) after first blocking with anti-CD16/32 antibody (1 ug/sample, BD FC block clone 24.G2). Viability was determined by staining with Fixable Viability Dye eFluor 780 (eBioscience). Samples were sorted on a BD FACS Aria III with a 70 μm nozzle, NK cells were identified as alive, NK1.1^+^ CD3^−^ and then immature as CD27^hi^CD11b^lo^ and mature CD27^lo^ CD11b^hi^ cells were sorted from the same animal. Total RNA was extracted from the cells using the Zymo DirectZol kit and stranded RNAseq libraries were prepared with Oligo dT pulldown according to the manufacturers protocol (Illumina). Libraries were sequenced on an Illumina HiSeq v4 to generate paired-end 75 bp reads that were aligned with STAR ^41^ to GRCm38 Mus musculus reference genome. Read counts were generated with HTseq ^42^ and differential expression analysis performed with DEseq2 ^43^.

### High throughput immunophenotyping

Cohorts of mixed sex mice with a range of genetically altered genotypes and WT mice were collected in batches of up to 48 in one collection with an age range of 12-16 weeks. In total it was aimed to collect data from 5 female and 5 male mice per allele. Mice were terminally anaesthetised with ketamine and xylazine and blood collected into EDTA coated tubes from the retro-orbital sinus. Whole blood or spleen single cell suspensions were incubated with titrated antibody cocktails to give saturating staining (details on Zendo 10.5281/zenodo.18879757). Spleen samples were first blocked with anti-CD16/32 antibody (1 ug/sample, BD FC block clone 24.G2) and DAPI (0.2 μg/ml) was added as a viability indicator. All samples were acquired on a 5 laser BD Fortessa that was setup using BD Cytometer Setup and Tracking beads, with application settings applied to ensure stability of the voltages. Compensation samples were prepared using Ultracomp eBeads (eBioscience) with compensation calculated using BD FACSDiva software. The data were analysed using a standardised template and gating strategy in FlowJo. Data was excluded if there were < 10,000 CD45+ singlet events.

### In vivo killing assay

Splenocytes from WT or B2M KO ^44^ (B6.129P2-*B2m^tm1Unc^*/DcrJ, RRID:IMSR_JAX:002087) mice were labelled with CFSE or CellTracker deep red and intravenously injected into recipient mice as indicated in the figure legend. After 90 min a 5 μl blood sample was collected into FACS buffer and red blood cell lysis was performed prior to acquisition on a flow cytometer (BD LSR II), transferred splenocytes were identified as CFSE^+^ or CellTracker deep red^+^ and the relative ratio between the two was determined and set at the baseline. This was repeated the following two days and the relative loss of the B2M KO relative to the WT determined.

### MCMV infection

For the majority of *in vivo* experiments, tissue culture-derived MCMV (pSM3fr-MCK-2fl BACmid) was grown and titred using 3T3 cells (ATCC, CRL-1658) with a carboxycellulose overlay. For salivary gland virus, MCMV Smith strain (ATCC) was prepared in the salivary glands of 3- to 4-week-old BALB/c mice and virus was purified over a sorbital gradient. Virus was passaged no less than 3 and no more than 5 times *in vivo*. Virus stocks and supernatant from homogenized organs isolated from MCMV-infected mice were titered for 6 days on a 3T3 cell monolayer with a carboxymethylcellulose overlay. Mice were infected with 1 x 10^6^ tissue culture-derived or 3 x 10^4^ PFU salivary gland-derived MCMV. Some mice were treated with 200μg anti-NK1.1 (clone PK136, BioXCell) or PBS control on days −2, 0 and +2 post infection Age- and sex-matched mice (males and females) aged 6-12 weeks were used.

### Flow cytometry

Splenocytes (1-2 × 10^6^) were stained with Zombie Aqua dye (Biolegend), incubated with Fc block (Biolegend), and stained for anti-CD3 (clone 145-2C11), anti-NK1.1 (clone PK136), anti-Ly49H (clone 3D10), anti-CD11b (clone M1/70), anti-CD27 (clone LG.7F9), anti-CD25 (clone PC61). Some cells were incubated for 5 hr with anti-CD107a (clone 1D4B), PMA and Ionomycin (both Sigma Aldrich) in the presence of monensin (for the final 4 hrs, Biolegend). Following surface staining, some cells were fixed and permeabilised with permeabilization buffer (ThermoFisher Scientific) and stained with anti-Granzyme B (clone GB11). All antibodies were purchased from Biolegend. At least 20,000 leukocytes were analysed using an Attune NxT Flow Cytometer (Thermo Fischer Scientific). Data were analysed using FlowJo software (v10.8.1, Tree Star).

### In vitro cytotoxicity and killing assays

For assessment of *in vitro* cytotoxicity, CD3^−^ cells were isolated (Miltenyi) and incubated for 48 hr with 1000 IU IL-2 (Proleukin). Cells were then stimulated +/− PMA and Ionomycin, IL-15 (30 ng/ml, Biolegend), IL-18 (60 ng/ml, Biolegend), or with P815 cells pre-incubated or not with 10 μg/ml anti-Ly49H or anti-NKG2D (both Biolegend) in the presence of anti-CD107a as described above. NK degranulation was expressed as % CD107a^+^ of NK1.1^+^CD3^−^ cells.

To measure cell killing, NK cells were isolated from spleens of naïve WT and *Sytl3*^−/−^mice (>78% purity) using negative selection (Miltenyi Biotec). BaF/3 control and BaF/3-m157 target cells (kind gifts from Professor Wayne Yokoyama, Washington University, USA) were stained with Cell Trace Violet (Invitrogen) and NK cells were co-incubated at various effector:target ratios as stated in the legend. After 4 hours, cells were stained with Annexin V-FITC according to the manufacturers’ instructions (Biolegend). Cell Trace Violet^+^ Annexin V^+^ cells were quantified using flow cytometry as described above.

### NK cell imaging

Purified NK cells were allowed to settle on anti-NK1.1 (PK136) coated slides for 10 min at 37°C in 150 μl of complete RPMI. Cells were then fixed and permeabilized with the addition of 150 μl of 8% PFA, 0.2% Triton X-100 in PBS at RT for 15 min. F-actin was stained with Alexa Fluor 647–labelled phalloidin (1:500 dilution in PBS; Invitrogen) and imaged by confocal microscopy (Leica-SP8-Inverted) with a 100x oil-immersion objective. Images were exported to ImageJ, and the percentage of cells with different distributions of F-actin were scored.

### Human orthologue

Mouse genes were mapped to the current gene name using Mouse Genome Informatics (https://www.informatics.jax.org) and human orthologues obtained from both Ensembl and Mouse Genome Informatics. LOUEF values were obtained from gnomAD v4.1.0 (https://gnomad.broadinstitute.org/downloads#v4-constraint). The gnomAD dataset was filtered for the mane select transcript and a single transcript per gene before the mouse orthologue mapping was performed. NK cell counts for the INTREPID cohort and controls were determined using BD Trucount tubes and a TBNK assay. Ethical approval was obtained from the North East - Newcastle & North Tyneside 2 Research Ethics Committee (REC 23/NE/0157). All participants provided written informed consent.

### Human knockout NK cell generation

gRNA sequences were obtained from the Vienna BioCenter website (https://www.vbc-score.org) and after addition of cloning homology arms were inserted into BbsI digested pKLV2-U6gRNA5(BbsI)-PGKpuro2AmCherry-W (a gift from Kosuke Yusa, Addgene plasmid # 67977) by Gibson Assembly (NEB HiFi DNA assembly master mix) according to the manufacturers protocol for single stranded DNA oligos. A safe targeting gRNA was selected from those previous annotated ^45^ and cloned as above. Virus-like particles (VLPs) were generated using pCMV-MMLVgag-3xNES-Cas9 (a gift from David Liu, Addgene plasmid # 181752), pBS-CMV-gagpol (a gift from Patrick Salmon, Addgene plasmid # 35614) and BaEV envelope (a gift from Daniel Hodson) as described ^46^ with the following modifications. In order to track uptake of the VLPs eGFP was fused to the C-terminal end of Cas9 with a GSS linker using pKLV2-U6gRNA5(gGFP)-PGKBFP2AGFP-W (a gift from Kosuke Yusa, Addgene plasmid # 67980) as a donor and PCR primers (FwdgaattcgagcccaagaagaagaggaaagtcggtagttccATGGTGAGCAAGGGCGAG, Rev cccataatttttggcagagggaaaaagatcTTACTTGTACAGCTCGTCCATG) and Phusion Plus PCR master mix. pCMV-MMLVgag-3xNES-Cas9 was linearised using PCR primers (Fwd GATCTTTTTCCCTCTGCC, Rev GACTTTCCTCTTCTTCTTG) with Phusion Plus PCR master mix. After DpnI digestion and clean-up of PCR products (NEB Monarch) Gibson reaction was setup with HiFi DNA assembly master mix (NEB) at a 2:1 ratio of insert:vector with a total of 0.2 pmol DNA. NEB stable high efficiency cells were transformed with the assembled DNA and positive clones checked via Sanger sequencing. The pCMV-MMLVgag-3xNES-Cas9-eGFP was site mutated to make the ‘v5’ derivative at C507V and pBS-CMV-gagpol was site mutated at Q226P using primers generated on NEBaseChanger and the Q5 Site-Directed Mutagenesis kit to improve efficiency as previous described ^47^. VLPs were produced in T175 Cell+ flasks (Sarstedt) seeding 17.5 x10^6^ HEK293T cells in 20 ml IMDM with 10% heat inactivated FBS six hours prior to transfection. Transfection was performed mixing 0.26 μg BaEV envelope, 7.88 μg pBS-CMV-gagpol(Q226P), 2.63 μg pCMV-MMLVgag(C507V)-3xNES-Cas9-eGFP and 10.27 μg gRNA vector in 200 μl OptiMEM media. Plus reagent was added at 1 μl per μg of DNA and Lipofectamine LTX added at a 2:1 ratio and incubated for 30 minutes at room temperature prior to addition to cells. After approximately 18 hours the media was replaced with 20 ml of fresh IMDM with 10% heat inactivated FBS. VLPs were harvested after 20-24 hours stored at 4°C and another 20 ml IMDM with 10% heat inactivated FBS. Two further rounds of VLPs were harvested and pooled prior to filtration through a 0.45 μM syringe filter. VLPs were concentrated 200x by addition of LentiX concentration solution (Takara) following the manufactures guidelines and the pellet was resuspended in RPMI 1640 with 10% heat inactivated FBS.

Peripheral blood NK cells were isolated from PBMCs (obtained from NHS blood and transplant non clinical issue, ethical approval REC 22/PR/1280 London Chelsea Research Ethics Committee, all subjects provide informed consent) using Human NK cell isolation kit (Miltenyi Biotec) and cultured in NK MACS media (Miltenyi Biotec) supplemented with 5% heat inactivated Human AB serum (Sigma), gentamycin (1 μg/ml, Gibco), 500 U/ml human IL-2 improved sequence (Miltenyi Biotec) and 10 U/ml human IL-15 (Miltenyi Biotec). After 6 - 7 days of expansion NK cells were harvested and VLPs were added (10 μl per 10^5^ cells) and centrifuged at 2000g for 90 min 32°C to enhance transduction. Approximately 24 hours later a small sample was collected to determine GFP fluorescence. NK cells were expanded for a further 7 days prior to functional assays and a small aliquot of cells taken to determine knockout efficiency via Sanger sequencing after PCR across the targeted region using Phire Tissue direct kit (ThermoFisher) and ICE online tool (EditCo).

### Human NK functional assays

NK cells were incubated in the presence of anti-CD107a B515 (Miltenyi Biotec) and BD GolgiStop (1 in 1500 final dilution) and either left unstimulated or stimulated with a 1:1 ratio of K562 cells (ATCC) or PMA (500 ng/ml) and Ionomycin (4 ng/ml) for 4 hours. Samples were washed with PBS and stained with anti-CD56 V423 and fixable/viability dye e506. After washing with FACS buffer cells were fixed with BD Cytofix/Cytoperm for 20 min at 4°C and washed twice with 1x BD Perm/Wash buffer. Intracellular staining with anti-IFNγ PE and anti-TNFα PE-Vio770 was performed overnight in 1x BD Perm/Wash buffer. Samples were washed twice prior to acquisition on a MACSQuant VYB instrument. Data was analysed in FlowJo and gated as singlets, live cells, CD56+ NK cells and positive for CD107a, IFNγ and TNFα.

K562 target cells were labelled with 2.5 μM TagIt violet dye (Biolegend) and co-cultured with NK cells at the indicated effector to target ratio for 3 hours. After washing with PBS cells were stained with fixable live/dead e780 dye and after washing with FACS buffer fixed with 2% paraformaldehyde. Samples were washed twice with FACS buffer prior to acquisition on a MACSQuant VYB instrument. Data was analysed in FlowJo and gated as target cells (SSC vs Violet) and the dead target cells (e780 vs FSC).

### Statistical analyses

For paired analysis (WT vs *Sytl3^−/−^* mice), statistical significance was determined in Prism using the Mann-Whitney U test or Student’s T-Test. Kruskall-Wallis was adopted for analysis of data derived from >2 groups of mice. *P* values are reported as follows: n.s.,>0.05; *, ≤0.05; **, ≤0.01; ***, ≤0.001; and ****, ≤0.0001. The primary immunophenotyping data was analysed with the PhenStat package in R ^8^ using the time as a fixed effect test to enable analysis of the multi-batch workflow using the concurrent controls. The p values were adjusted for multiple testing with the Benjamini & Hochberg correction per tissue and the z score of the effect size derived for each cell type on the basis of sex to enable application of an effect size cutoff. Human functional NK data was analysed using a two-way ANOVA with correction for multiple testing with a post-hoc Dunnett test in Prism.

---

**Figure S1.**
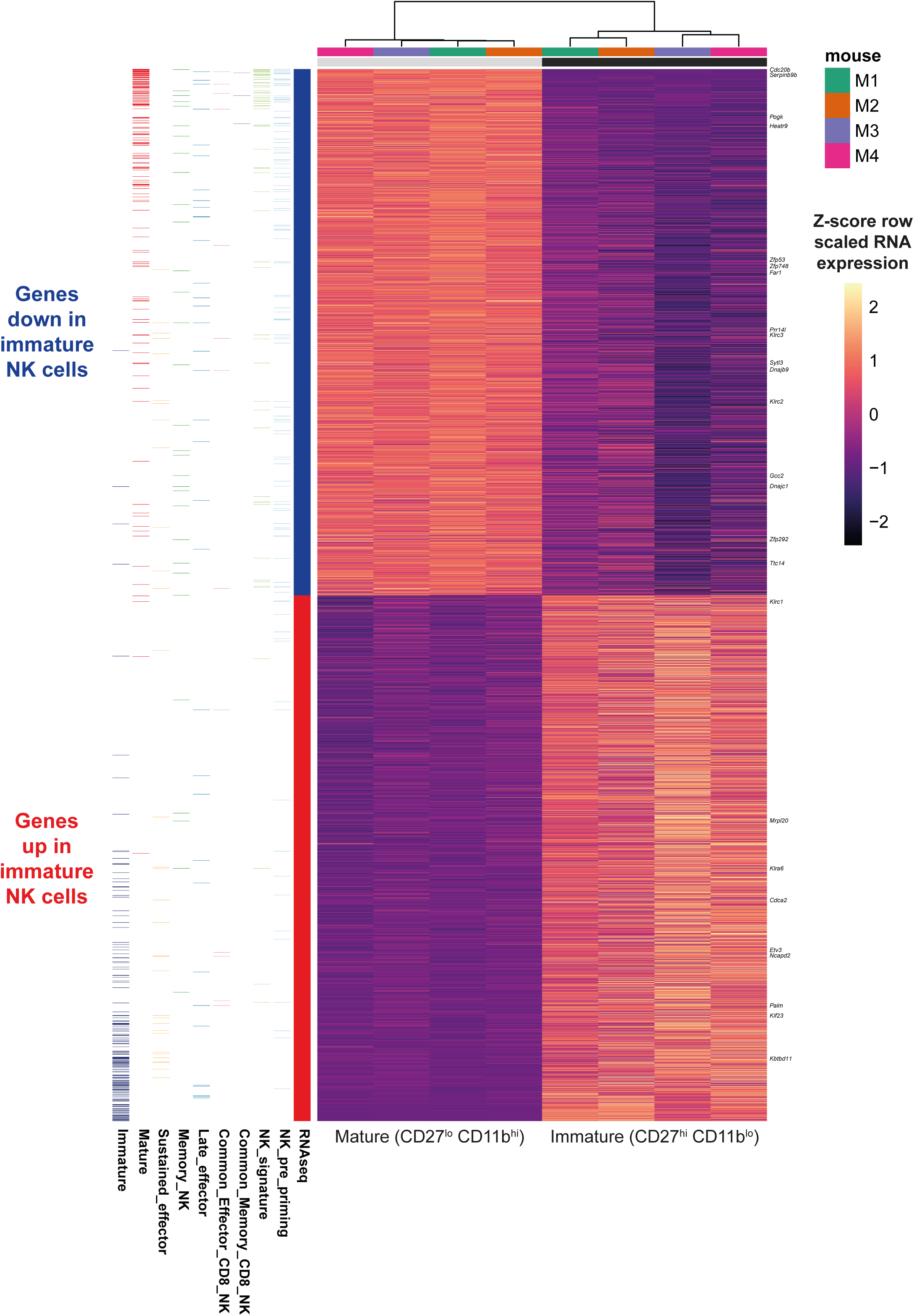
Heatmap of significantly differentially expressed genes between immature and mature NK cells. RNAseq normalised scaled read counts for genes differentially expressed (DEseq2 Padj < 0.05) between immature (CD27^hi^ CD11b^lo^) and mature (CD27^lo^ CD11b^hi^) sorted splenic NK cells. Genes analysed as mouse KO are indicated and the gene assignment from previous microarray experiments immature and mature NK ^4^ or effector functions ^7^.

**Figure S2.**
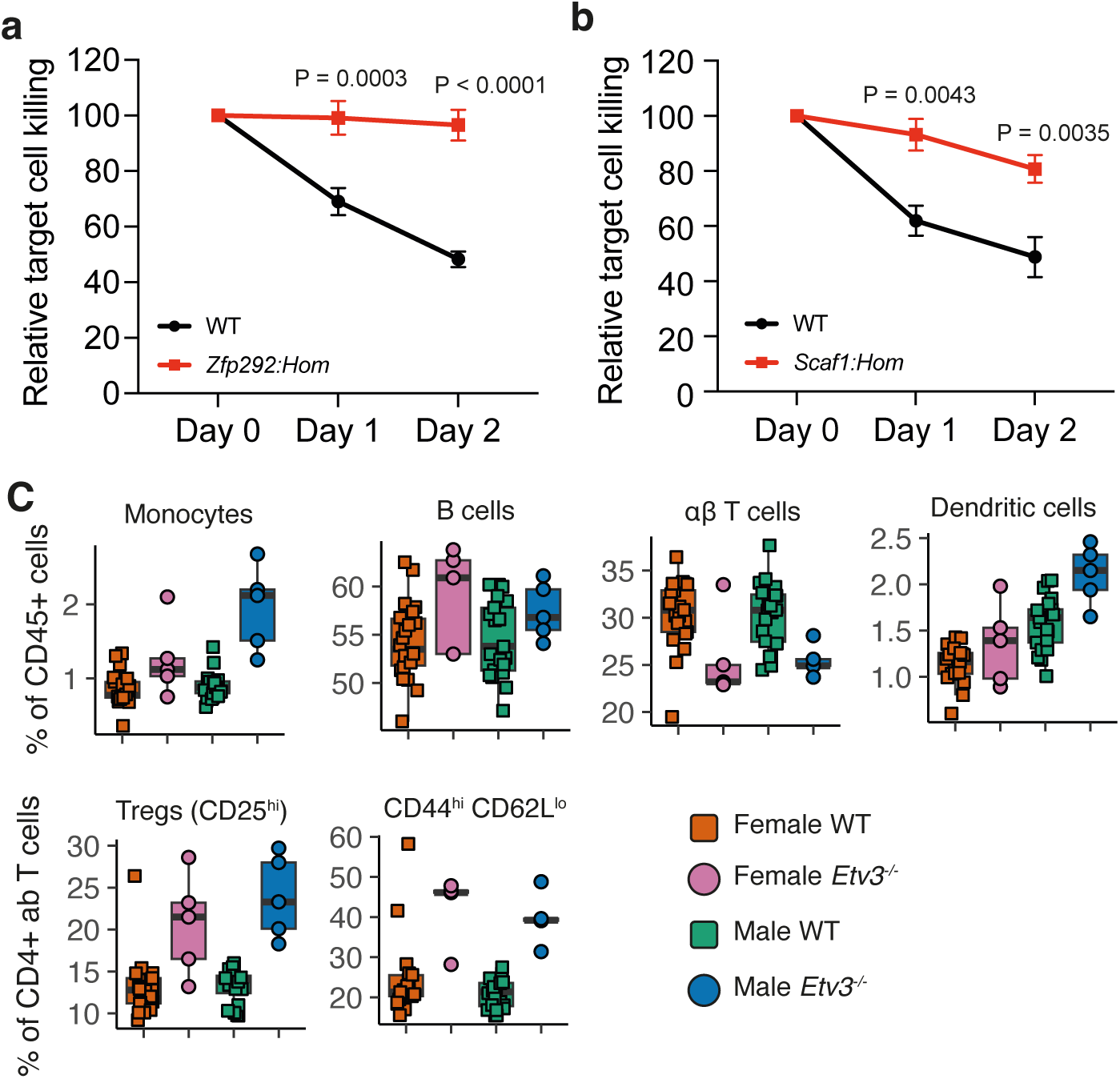
Impaired in vivo killing and Etv3 baseline immunophenotyping. **(a-b)** *In vivo* killing of B2M WT vs B2M KO splenocytes in *Zfp292^−/−^* or *Scaf1^−/−^* mice. Symbol is mean ± SEM with n = 7 WT and *Zfp292^−/−^* and n = 10 WT and n = 5 *Scaf1^−/−^.* Statistical analysis was performed with a 2-way repeated measures ANOVA with an uncorrected Fisher’s LSD post-hoc test for each timepoint. **(c)** Baseline immunophenotyping data for *Etv3^−/−^* mice, each symbol represents an individual animal with the gene and parameter listed above the plot. All reported are significant findings with padj < 0.05.

**Figure S3.**
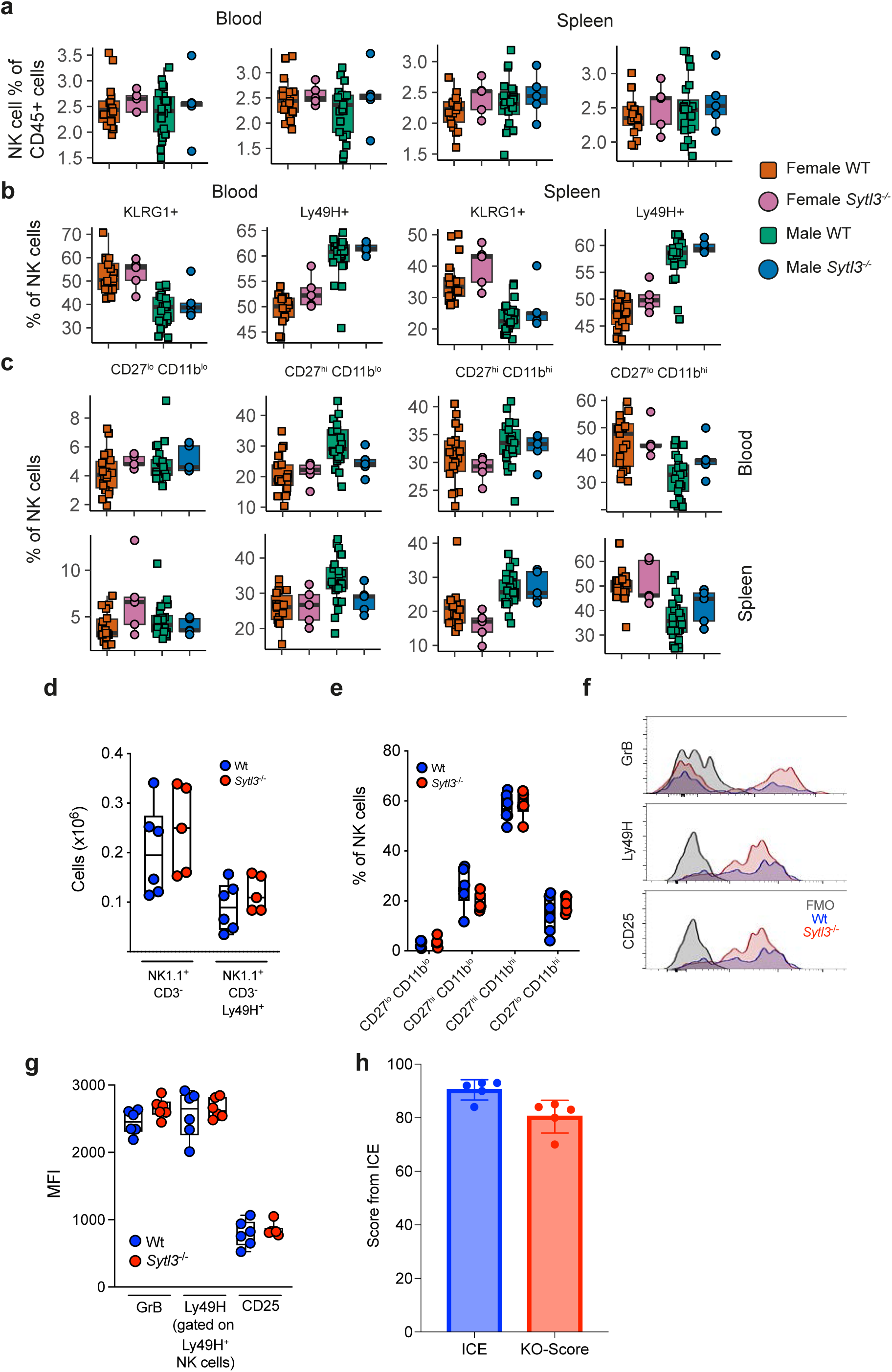
Sytl3 does not impact NK cell development or phenotype. (**a-c)** NK baseline immunophenotyping from *Sylt3*^−/−^ mice. **(a)** NK cells are expressed as % of CD45+ live cells from blood and spleen. **(b-c)** NK phenotypic subsets are % of NK cells with the marker or subpopulation indicated above the graph. Each symbol represents an individual animal. NK cell numbers **(d)** and NK cell developmental subsets on the basis of CD27 and CD11b **(e)** in spleen after MCMV infection. **(f & g)** Expression of Granzyme B, Ly49H and CD25 on NK cells in the spleen after MCMV infection. All MCMV infection experiments were performed at least twice. **(h)** Assessment of KO of Human NK cells using SYTL3_g4 determined by EditCo ICE tool, ICE indicates the total indel fraction and KO are predicted deleterious indels that would result in a functional KO. Due to the genomic sequence around the location of SYTL3_g1 it was not possible to generate a product that would yield a Sanger sequencing trace.

## Acknowledgments

We acknowledge the services provided by the Cambridge Human Immunophenotyping Platform, the NIHR Cambridge BRC Cell Phenotyping Hub, as well as the Cytometry Core Facility, DNA pipelines and Informatics at the Wellcome Sanger institute (supported by Wellcome Grant 206194). We are grateful for the facilities and support provided by the Wellcome Sanger Institute Research Support facility, University of Cambridge University Biological Services Unit and Cardiff University.

AOS was supported by a package from Wellcome Sanger Institute and Wellcome Trust (206194). IH was supported by the MRC (MR/X00922X/1, APP40526), Wellcome Trust (207503/Z/17/Z) and Kidney Research UK/Kidney Wales (ST_003_20220705). SC, KH, CBR, CB, AA, BAH, VI, LvdW and DJA were supported by the Wellcome Trust (206194). EP was supported by a Cardiff University School of Medicine PhD Studentship. ECYW was supported by the MRC (MR/P001602/1, MR/V000489/1) and Kidney Research UK (RP_008_20210729). PAL and MCC were supported by the Wellcome Trust (219506/Z/19/Z). DMD was supported by the MRC (MR/W031698/1). RH was supported by a Wellcome Trust PhD studentship.

## Author contributions

Conceptualization; IH and AOS

Validation; EP, SC, KH, LvdW, AOS

Formal analysis; SS, VI, MCC, DD, IH, AOS

Investigation; EP, SC, RH, SS, MC, LC, KH, CBR, CB, AA, BAH, LvdW, MAO

Resources; PAL, DJA, MCC, IH and AOS

Supervision; DJA, DD, ECYW, IH and AOS

Data Curation; EP, MM, PAL, MCC, IH and AOS

Funding acquisition; DJA, ECYW, DD, IH, and AOS

Visualization; IH and AOS

Writing – original draft preparation; IH and AOS

Writing – review and editing; all authors

## Data availability

The RNA sequencing data is available from the European Nucleotide Archive under accession (TBC, release in progress). The full immunophenotyping data, statistical analysis, metadata and immunophenotyping antibody details are available on Zendo 10.5281/zenodo.18879757.

## Code availability

Code available upon request to the corresponding author (AOS).

## INTREPID consortium authors

Katarina Pajerska^1^, Rachael Bashford-Rogers^2^, Siobhan Burns^3,4^, Christoph Hess^1^, Paul Lyons^1^, Kenneth Smith^1^, Ernest Turro^5^, Chris Wallace^1^, Sarah Wordsworth^6^, and Matthew Cook^1^

^1^ Cambridge Institute for Therapeutic Immunology and Infectious Diseases, Department of Medicine, University of Cambridge

^2^ Department of Biochemistry, University of Oxford

^3^ University College London Institute of Immunity and Transplantation

^4^ Department of Immunology, Royal Free London NHS Foundation Trust

^5^ Icahn School of Medicine at Mount Sinai, NY

^6^ Nuffield Department of Population Health, University of Oxford

